# Guard Cell Chloroplast Remodeling Under Salt Stress Correlates with Ion Enrichment in Faba Bean

**DOI:** 10.64898/2026.08.11.744119

**Authors:** Xudong Zhang, Guanghui Wei, Marc Welzer, Christian Zörb

## Abstract

Salinity stress alters cellular ion homeostasis and photosynthetic activity, yet how guard cell chloroplast architecture contributes to salt adaptation remains poorly understood. Here, we investigated salt-induced chloroplast remodeling in guard cells of two faba bean genotypes, Fuego and Scoop, by integrating 3D chloroplast imaging, ion enrichment analysis and photosynthetic measurements. Salt stress induced distinct genotype-dependent changes in chloroplast morphology, with Fuego exhibiting pronounced chloroplast enlargement and reduced surface area-to-volume ratios under Na₂SO₄ and high NaCl, whereas Scoop showed treatment-dependent remodeling with larger chloroplasts under low NaCl and higher surface area-to-volume ratios under Na₂SO₄ and CaCl₂. These structural responses were associated with differential Na and Cl partitioning at the stomatal complex surface. In Fuego, chloroplast size was negatively associated with photosynthetic rate, whereas Scoop showed positive relationships between chloroplast size and photosynthetic performance. Multivariate analysis further revealed coordinated associations among chloroplast architecture, ion enrichment and photosynthesis that distinguished the two genotypes under salinity. Our findings demonstrate that guard cell chloroplast remodeling is closely associated with genotype-specific salt responses and local ion partitioning. Integrating organelle structural plasticity with local ion homeostasis provides a spatially resolved perspective on the cellular basis of genotype-dependent salinity adaptation.

## 1 Introduction

Chloroplasts are highly dynamic organelles that continuously adjust their structure and function in response to environmental fluctuations (Kirchhoff, 2019; Schwenkert et al., 2022). Beyond carbon fixation (Shi et al., 2024), chloroplasts integrate energy metabolism, redox homeostasis and stress signaling, enabling plants to reprogram cellular metabolism under adverse conditions (Collado-Arenal et al., 2024; Jarvis and López-Juez, 2013). Changes in chloroplast morphology, including alterations in size, shape and internal organization, have been documented under a wide range of abiotic stresses and are considered important components of plant acclimation (Oi et al., 2020; Yamane et al., 2018). However, the functional significance of stress-induced chloroplast remodeling remains poorly understood, particularly in specialized cell types that directly regulate plant responses to the environment.

Guard cells represent a unique photosynthetic cell type because they integrate light perception, metabolism and ion transport to regulate stomatal aperture (Lawson and Matthews, 2020). Unlike mesophyll chloroplasts, guard cell chloroplasts directly couple photosynthetic metabolism with stomatal function and differ in morphology, metabolic activity and physiological roles (da Silva et al., 2025; Lawson, 2009; Zeiger et al., 2002). During stomatal movements, guard cell chloroplasts contribute to the synthesis and turnover of metabolites, including starch, which are closely linked to osmotic regulation and aperture control (Dang et al., 2024; Granot and Kelly, 2019; Lim et al., 2022; Santelia and Lunn, 2017). Despite their functional importance, how guard cell chloroplast architecture responds to environmental stress and whether these structural changes contribute to stomatal adaptation remain largely unexplored.

Salinity provides a particularly relevant context for investigating guard cell chloroplast plasticity because it simultaneously imposes osmotic and elemental constraints (Hedrich and Shabala, 2018; Zörb et al., 2022). Excess Na and Cl accumulation disrupts cellular homeostasis, while elemental compartmentation influences photosynthesis, osmotic adjustment and stress signaling (Munns and Tester, 2008; Zörb et al., 2019). Previous studies have shown that salt stress promotes chloroplast starch accumulation and ultrastructural remodeling, including thylakoid reorganization and changes in stromal organization (Naeem et al., 2012; Thalmann and Santelia, 2017; Wang et al., 2013). Recent imaging studies further revealed the spatial co-localization of salt elements and starch granules within chloroplasts, suggesting a close coordination between elemental homeostasis and carbon metabolism during salinity adaptation (Kanai et al., 2007; Noda et al., 2023). Nevertheless, whether such remodeling occurs in guard cell chloroplasts and how it is associated with local elemental partitioning and photosynthetic performance remain unknown.

Here, we investigated salt-induced guard cell chloroplast remodeling in two faba bean genotypes with contrasting physiological responses. We hypothesized that (*i*) salinity triggers genotype-dependent changes in guard cell chloroplast architecture; (*ii*) these structural responses are associated with differential Na and Cl partitioning within stomatal complexes; and (*iii*) the coordination between chloroplast remodeling and local elemental homeostasis contributes to genotype-specific photosynthetic performance under salt stress. By integrating 3D chloroplast imaging, stomatal-complex elemental analysis and gas-exchange measurements, we reveal how local elemental partitioning is associated with chloroplast remodeling and photosynthetic performance under salinity.

## 2 Materials and methods

### 2.1 Plant cultivation

Two contrasting faba bean (*Vicia faba* L.) genotypes, Fuego and Scoop (Norddeutsche Pflanzenzucht Hans-Georg Lembke KG, Hohenlieth, Germany), were cultivated hydroponically in a climate cabinet (WEISS HGC1014, Heuchelheim, Germany) under a 14 h light/10 h dark photoperiod, 22/18 °C day/night temperatures, approximately 80–60% relative humidity, and 300 μmol photons m⁻² s⁻¹ at shoot level. Seeds were initially soaked in aerated 0.5 mM CaSO₄ solution for 24 h at room temperature and then germinated in moistened quartz sand. Twelve days after germination, seedlings were transferred to plastic pots containing one-quarter-strength aerated nutrient solution. Nutrient strength was gradually increased to full strength by sequential additions of one-half, three-quarter and full-strength solutions over nine days. The full-strength nutrient solution contained 0.1 mM KH₂PO₄, 1.0 mM K₂SO₄, 2.0 mM Ca(NO₃)₂, 0.5 mM MgSO₄, 0.00464% (w/v) Sequestren (Ciba Geigy, Basel, Switzerland), 10 μM NaCl, 10 μM H₃BO₃, 2.0 μM MnSO₄, 0.5 μM ZnSO₄, 0.2 μM CuSO₄, 0.1 μM CoCl₂ and 0.05 μM (NH₄)₆Mo₇O₂₄.

After acclimation to full-strength nutrient solution, plants were subjected to five salt treatments: control (1 mM NaCl), 50 mM NaCl, 100 mM NaCl, 50 mM Na₂SO₄ and 50 mM CaCl₂. Salt concentrations were increased gradually over three days by adding one-third of the final concentration each day to minimize osmotic shock. Plants were harvested eight days after full salt exposure. Gas exchange measurements were performed on the fifth fully expanded leaf between 4 and 6 h after the onset of the light period for seven consecutive days following salt exposure. For chloroplast imaging, leaf segments from the fifth fully expanded leaf were immediately immersed in liquid nitrogen and stored at −80 °C until confocal microscopy analysis. Epidermal strips from the fifth fully expanded leaf used for gas exchange measurements were collected, immediately cryopreserved in liquid nitrogen and stored until cryo-SEM-EDX analysis to preserve native stomatal complex elemental distribution. Remaining leaf tissues were frozen, ground and freeze-dried for ICP-OES analysis. Each treatment consisted of three pots with three plants per pot.

### 2.2 Gas exchange measurements

Gas exchange measurements were performed using a portable photosynthesis system (GFS-3000, Heinz Walz GmbH, Effeltrich, Germany). Net photosynthetic rate (A) was used as the primary physiological indicator of photosynthetic performance. Measurements were conducted on the fifth fully expanded leaf under controlled conditions with a cuvette area of 2.5 cm², 400 ppm CO₂, 15,000 ppm H₂O, 25 °C air temperature, 1000 μmol m⁻² s⁻¹ photosynthetically active radiation (PAR), and a gas flow rate of 750 μmol s⁻¹. Additional gas exchange parameters, including transpiration rate (E) and stomatal conductance (gs), were recorded. At least five measurement spots were obtained from five independent plants per treatment.

### 2.3 Relative ion enrichment patterns in stomatal complex surface regions

Leaf ion concentrations measured by ICP-OES and stomatal complex (StC) elemental signals obtained by cryo-SEM-EDX were used to quantify relative ion partitioning between bulk leaf tissues and stomatal complex surface regions (Supplementary Methods S1 leaf ionome and S2 StC surface element and Figure S1). The enrichment index was calculated as the log₁₀ ratio between normalized StC and leaf ion responses:

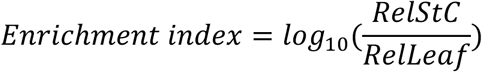

where ion responses were normalized to the corresponding genotype-specific control. A small pseudocount (ε), defined as half of the minimum non-zero value across all samples, was applied before ratio calculation and log₁₀ transformation to avoid undefined values caused by zero or near-zero measurements (Lambert et al., 1991; Martín-Fernández et al., 2003). The ε value was defined as half of the minimum non-zero value across all samples. Sensitivity analysis using alternative ε values (0.5×-2×) confirmed consistent enrichment patterns across genotypes and treatments (Figure S2).

Positive enrichment index values indicate relative ion enrichment in stomatal complex surface regions compared with bulk leaf tissues, whereas negative values indicate relative depletion.

### 2.4 3D chloroplast imaging and morphological analysis

Fully expanded leaves, used for gas exchange monitoring, were harvested from independent plants within each treatment and stored at −80 °C. Frozen leaves were transferred directly into ice-cold 1× PBS (pH 7.3) to avoid air-thawing, then thawed gradually while fully submerged on ice for 10–15 min to allow gentle buffer infiltration while preserving tissue integrity. Leaves from each treatment were pooled, and stomatal complexes were randomly selected for confocal imaging and quantitative analysis.

Confocal images were acquired using a Zeiss LSM900 confocal microscope (Zeiss Microscopy GmbH) equipped with a Plan-Apochromat 40×/1.2 Imm Corr DIC M27 objective. Chloroplasts were visualized based on chlorophyll autofluorescence excited with the 640 nm laser line, while transmitted light images were obtained using the ESID detector. Z-stack images were collected from target individual stomatal complexes for 3D reconstruction.

Images were segmented and analysed using ArivisPro (Zeiss Microscopy GmbH) with a deep-learning model trained in ArivisCloud (Zeiss Microscopy GmbH). The model was trained by manual annotation of one representative image per experimental condition and validated using three randomly selected planes from four independent images excluded from training. Segmentation performance was evaluated against manual annotations using an Intersection over Union (IoU) threshold of 0.5, with F1-scores, precision and recall values above 0.5 for both chloroplast and guard cell segmentation. The validated model was subsequently used for stomatal and chloroplast segmentation, and the “compartments” function was applied to assign chloroplasts to individual guard cells. Chloroplast number, volume, surface area, surface area-to-volume ratio and sphericity were automatically extracted during batch analysis. All reconstructed images were manually inspected after batch processing to verify segmentation quality.

Because chlorophyll autofluorescence originates exclusively from chloroplasts, the 3D reconstruction workflow was optimized for accurate chloroplast visualization rather than quantitative reconstruction of complete guard cell morphology. Therefore, guard cell structural parameters were not used for biological interpretation. Guard cell reconstructions were retained only as a 3D framework to define chloroplast localization within guard cells.

A size-based quality control procedure was applied to remove potential segmentation artifacts. Different volume thresholds were evaluated, resulting in removal of 0.7–10.3% of detected objects depending on the selected cutoff (Supplementary Table S1). A conservative threshold of 0.2 µm³ was selected because it removed small isolated objects while retaining more than 90% of detected chloroplast objects, thereby minimizing exclusion of potentially biologically relevant chloroplast structures. All downstream analyses were performed using chloroplast objects above this threshold.

### 2.5 Principal component analysis

Principal component analysis (PCA) was performed to visualize genotype- and treatment-associated patterns in multivariate responses to salt stress. The analysis integrated chloroplast morphological traits, gas exchange parameters, leaf ion concentrations and stomatal complex ion enrichment indices to evaluate whether the two faba bean genotypes exhibited distinct response strategies associated with chloroplast remodeling and ion partitioning under different salt treatments.

### 2.6 Trait correlation analysis

Correlation analysis was performed as a complementary approach to PCA to examine pairwise relationships among physiological, ionic and chloroplast morphological traits under salt stress. The same set of parameters included in PCA, including chloroplast morphological traits, gas exchange parameters, leaf ion concentrations and stomatal complex ion enrichment indices, were integrated to generate genotype-specific correlation matrices. Correlation patterns were analysed separately for the two faba bean genotypes to identify genotype-dependent associations among chloroplast remodeling, ion partitioning and physiological responses.

### 2.7 Statistical analysis

Data were analysed using two-way analysis of variance (ANOVA) to assess the effects of genotype, treatment and their interaction (genotype × treatment). Multiple comparisons among treatments within each genotype were performed using Fisher’s least significant difference (LSD) test implemented in the agricolae package in R, whereas pairwise comparisons between genotypes within the same treatment were conducted using Welch’s t-test. Gas exchange measurements were based on five biological replicates per treatment. Ion enrichment indices were calculated as log₁₀ ratios between normalized stomatal complex surface elemental signals and corresponding whole-leaf ion concentrations, with at least 10 stomatal complexes analysed per treatment. Confocal microscopy analyses for chloroplast visualization and 3D reconstruction were performed using six stomatal complexes per treatment collected from pooled samples of five individual plants. Multivariate analyses, including PCA and correlation analysis, were performed in R using the prcomp function, ggplot2 and pheatmap packages.

## 3 Results

### 3.1 Guard cell chloroplast remodeling under salt stress in faba bean

To characterize salt-induced structural changes within stomatal complexes, 3D confocal reconstructions were performed to quantify guard cell chloroplast morphology in Fuego and Scoop (Fig. 1). Given the variable orientation and positioning of individual chloroplasts within guard cells, representative images illustrate the range of chloroplast configurations, whereas quantitative analyses based on 3D reconstructions were used to evaluate treatment effects. Two-way ANOVA revealed significant treatment × genotype interactions for chloroplast surface area and volume (P < 0.01 for both traits). Visually, chloroplasts under control conditions typically exhibited elongated, lenticular structures in both genotypes (Fig. 1a,f). Under salt treatments, chloroplast morphology shifted towards enlarged and more rounded structures, particularly in Fuego under Na₂SO₄ and high NaCl treatments (Fig. 1c,d) and in Scoop under low NaCl (Fig. 1g). In contrast, chloroplasts under CaCl₂ treatment appeared less swollen in both genotypes (Fig. 1e,i), consistent with the lower chloroplast dimensions observed in quantitative analyses.

**Figure 1.**
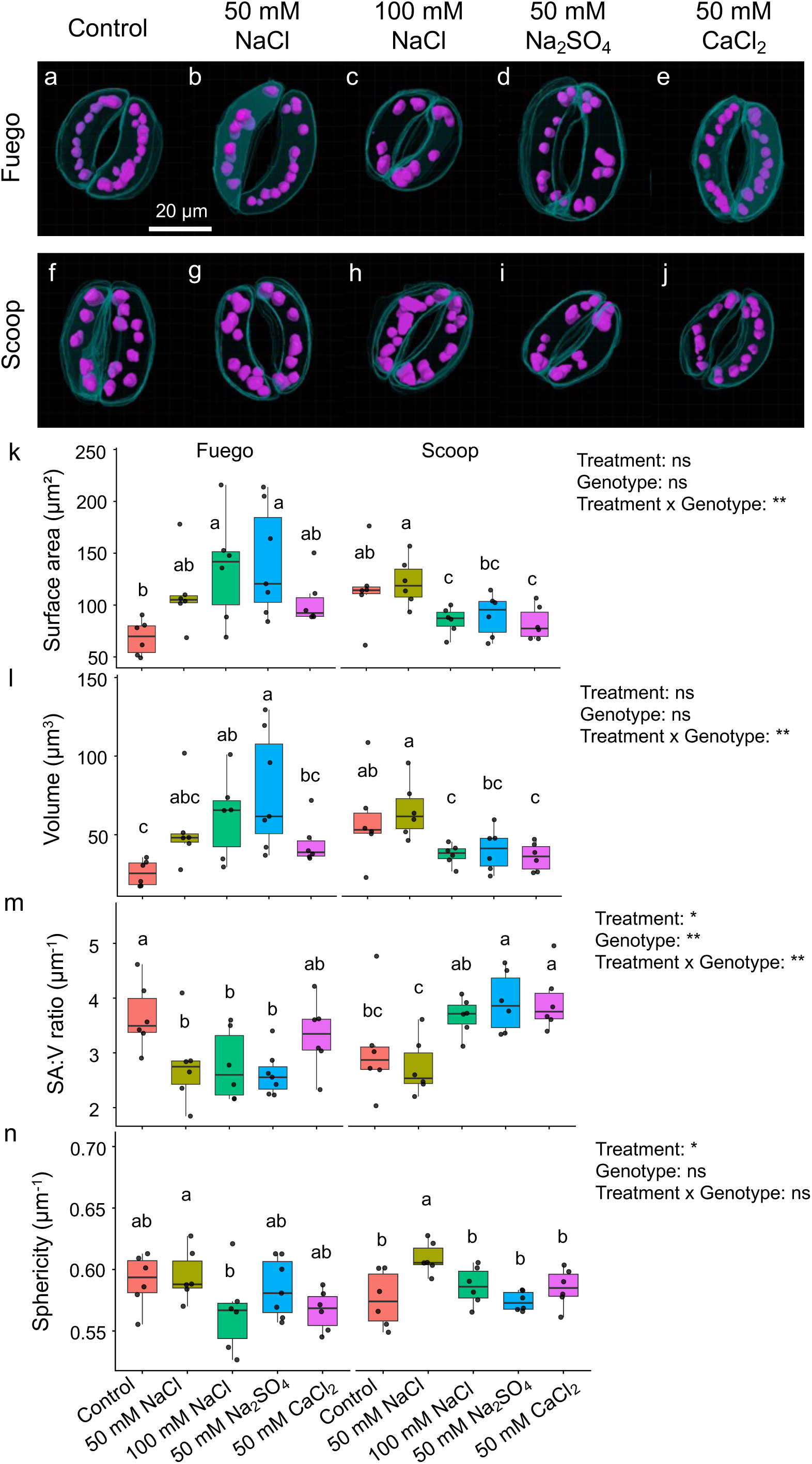
Guard cell chloroplast remodeling under salt stress in faba bean. a–j, Representative 3D reconstructions of chloroplasts (magenta) within guard cells (cyan) of the faba bean genotypes Fuego (a–e) and Scoop (f–j) under control and salt treatments. k–n, Quantification of chloroplast surface area (k), volume (l), surface area-to-volume ratio (SA:V ratio) (m) and sphericity (n) in guard cells of both genotypes. Data are presented as means ± s.e. (n = 6 stomatal complexes per treatment). Statistical significance was assessed using two-way ANOVA (ns, not significant; *, P < 0.05; **, P < 0.01), followed by Fisher’s LSD test for multiple comparisons. Different lowercase letters indicate significant differences among treatments within the same genotype (P < 0.05).

Quantitative analyses of chloroplast surface area and volume showed contrasting genotype-dependent responses to salt treatments (Fig. 1k,l). In Fuego, chloroplast surface area and volume increased under 100 mM NaCl and Na₂SO₄ relative to the control, reaching 141.8 µm² and 77.8 µm³, respectively, under Na₂SO₄ compared with 68.6 µm² and 25.7 µm³ under control conditions. The 50 mM NaCl and CaCl₂ treatments showed intermediate chloroplast dimensions that did not differ significantly from the control. In Scoop, chloroplast dimensions were highest under 50 mM NaCl, although they did not differ significantly from the control, whereas 100 mM NaCl and CaCl₂ resulted in significantly smaller chloroplasts. Na₂SO₄ produced intermediate chloroplast dimensions. Genotypic comparisons further revealed treatment-specific differences in chloroplast remodeling. Under control conditions, Scoop exhibited larger chloroplasts than Fuego, with surface area and volume of 115.7 µm² and 59.2 µm³ compared with 68.6 µm² and 25.7 µm³, respectively (P < 0.05). In contrast, under Na₂SO₄ treatment, Fuego showed greater chloroplast expansion than Scoop, with surface area and volume reaching 141.8 µm² and 77.8 µm³ compared with 90.2 µm² and 40.4 µm³, respectively (P < 0.05). No significant genotypic differences were detected under NaCl or CaCl₂ treatments (Table S1).

The surface area-to-volume (SA:V) ratio showed distinct genotype-dependent responses (Fig. 1m). In Fuego, the SA:V ratio decreased from 3.67 under control conditions to approximately 2.6–2.8 under NaCl and Na₂SO₄ treatments, with significantly lower values under 50 mM NaCl, 100 mM NaCl and Na₂SO₄ than under control conditions. In contrast, Scoop showed higher SA:V ratios under Na₂SO₄ and CaCl₂, reaching 3.93 and 3.94, respectively, whereas the lowest value was observed under 50 mM NaCl. Significant genotypic differences were detected under 100 mM NaCl and Na₂SO₄, where Scoop showed higher SA:V ratios than Fuego (Table S1).

Chloroplast sphericity was significantly affected by treatment but not genotype or the treatment × genotype interaction (Fig. 1n). Both genotypes showed their highest sphericity under 50 mM NaCl, reaching 0.595 in Fuego and 0.609 in Scoop. In Scoop, sphericity under 50 mM NaCl was significantly higher than under all other treatments, whereas in Fuego it was significantly higher than under 100 mM NaCl but did not differ significantly from control, Na₂SO₄ or CaCl₂. No significant genotypic differences were detected across treatments (Table S1).

### 3.2 Photosynthetic rate and ion enrichment indices in guard cells of two faba bean genotypes under salt stress

To evaluate the relationship between photosynthetic performance and ion partitioning within stomatal complexes, net photosynthetic rate (A) and Na and Cl enrichment indices were quantified in Fuego and Scoop across different salt treatments (Table 1). Two-way ANOVA revealed significant effects of genotype, treatment and their interaction for all three parameters (P < 0.001).

**Table 1.** Photosynthetic rate and ion enrichment indices in guard cells of two faba bean genotypes under salt stress. Values are presented as means ± s.e. For net photosynthetic rate (A), n = 5 biological replicates (individual plants). For Na⁺ and Cl⁻ enrichment indices, at least 10 stomatal complexes were analysed per treatment from five independent leaf samples. The enrichment index quantifies the relative accumulation or depletion of ions in the stomatal complex relative to the whole leaf. Different lowercase letters within the same genotype indicate significant differences among treatments (Fisher’s LSD test, P < 0.05). Asterisks indicate significant differences between genotypes within the same treatment (Welch’s t-test). *, ** and *** denote P < 0.05, P < 0.01 and P < 0.001, respectively. Asterisks are placed next to the value of the genotype with the higher mean. Two-way ANOVA revealed significant effects of treatment, genotype and their interaction for all three parameters (A, Na enrichment index and Cl enrichment index; all P < 0.001).

| Genotype | Treatment | A ($\mu\text{mol CO}_2 \text{ m}^{-2} \text{ s}^{-1}$) | Na enrichment index | Cl enrichment index |
| --- | --- | --- | --- | --- |
| Fuego | Control | 8.02 ± 0.29 a | -0.58 ± 0.00 b | -1.90 ± 0.00 e |
| Fuego | 50 mM NaCl | 6.89 ± 0.31 a* | -1.46 ± 0.04 c | -1.02 ± 0.00 c |
| Fuego | 100 mM NaCl | 4.67 ± 0.28 b | -1.48 ± 0.09 c | -1.32 ± 0.00 d |
| Fuego | 50 mM Na<sub>2</sub>SO<sub>4</sub> | 4.25 ± 0.28 b | -2.35 ± 0.00 d | -0.44 ± 0.00 a |
| Fuego | 50 mM CaCl<sub>2</sub> | 6.70 ± 0.31 a** | -0.23 ± 0.00 a | -0.87 ± 0.06 b |
| Scoop | Control | 8.63 ± 0.37 a | -1.06 ± 0.00 d | -1.51 ± 0.00 d |
| Scoop | 50 mM NaCl | 6.03 ± 0.23 b | -0.72 ± 0.10 c*** | -1.27 ± 0.00 c |
| Scoop | 100 mM NaCl | 4.05 ± 0.22 c | -0.55 ± 0.11 b*** | -1.49 ± 0.00 d |
| Scoop | 50 mM Na<sub>2</sub>SO<sub>4</sub> | 4.92 ± 0.27 bc | -1.71 ± 0.00 e | -0.50 ± 0.00 a |
| Scoop | 50 mM CaCl<sub>2</sub> | 5.26 ± 0.33 bc | 0.25 ± 0.00 a | -0.66 ± 0.01 b** |

Net photosynthetic rate showed genotype- and treatment-dependent responses to salinity. In Fuego, A remained comparable to the control under low NaCl and CaCl₂ treatments, with values of 6.89 and 6.70 µmol m⁻² s⁻¹, respectively, but decreased significantly under high NaCl and Na₂SO₄ to 4.67 and 4.25 µmol m⁻² s⁻¹. In Scoop, A decreased progressively with increasing NaCl concentration, from 8.63 µmol m⁻² s⁻¹ under control conditions to 6.03 and 4.05 µmol m⁻² s⁻¹ under low and high NaCl treatments, respectively. Fuego exhibited higher A than Scoop under low NaCl and CaCl₂, whereas no significant genotypic differences were detected under control, high NaCl or Na₂SO₄ treatments.

Na enrichment indices revealed distinct ion partitioning patterns between genotypes. In Fuego, Na was depleted from stomatal complexes under all salt treatments, with the strongest depletion under Na₂SO₄ (−2.35) and high NaCl (−1.48). In Scoop, Na depletion was weaker under both NaCl treatments, with enrichment indices of −0.72 and −0.55 under low and high NaCl, respectively; both values were significantly higher than those in Fuego under the corresponding treatments (P < 0.001). Under Na₂SO₄, Fuego showed numerically stronger Na depletion than Scoop (−2.35 versus −1.71), although the genotypic difference was not significant. Under CaCl₂, Scoop showed slight Na enrichment (0.25), whereas Fuego remained slightly depleted (−0.23), with no significant genotypic difference.

Cl enrichment showed a distinct treatment-dependent pattern in both genotypes. Na₂SO₄ resulted in the least negative Cl enrichment indices in Fuego and Scoop (−0.44 and −0.50, respectively), whereas control conditions showed the strongest Cl depletion (−1.90 and −1.51, respectively). Under CaCl₂, Scoop exhibited a significantly higher Cl enrichment index than Fuego (−0.66 versus −0.87, P < 0.01). NaCl treatments generally resulted in stronger Cl depletion than Na₂SO₄ in both genotypes.

### 3.3 Multivariate integration of chloroplast morphology, ion enrichment and photosynthetic performance in faba bean genotypes

To integrate chloroplast morphology, guard cell ion partitioning and photosynthetic performance under salt stress, principal component analysis was performed using chloroplast structural traits, net photosynthetic rate (A), and Na and Cl enrichment indices (Fig. 2). The first two principal components explained 72.4 % of the total variation, with PC1 and PC2 accounting for 49.4 % and 23.0 %, respectively.

**Figure 2.**
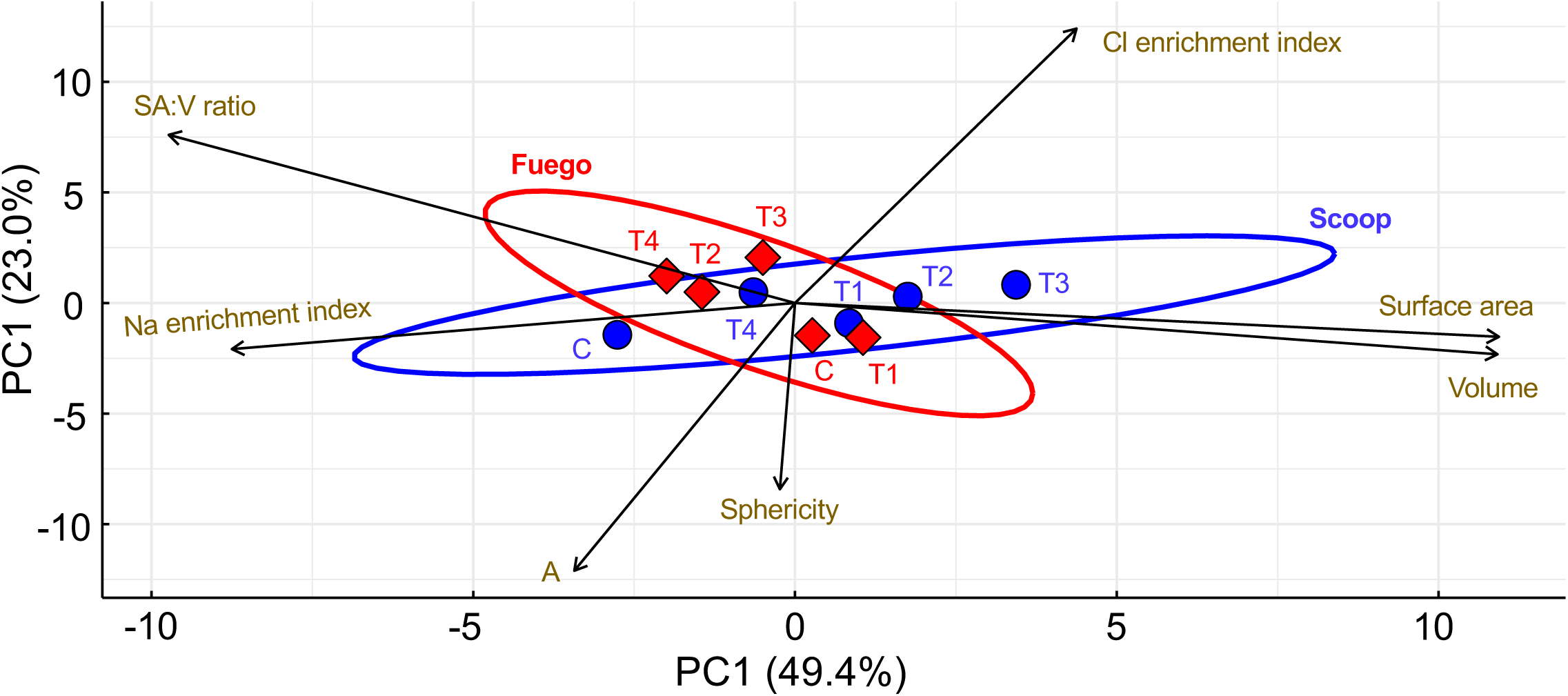
Principal component analysis of chloroplast morphology, ion enrichment and photosynthetic performance in faba bean genotypes. The PCA biplot shows multivariate relationships among chloroplast morphological traits (surface area, volume, surface area-to-volume ratio (SA:V) and sphericity), net photosynthetic rate (A), and Na and Cl enrichment indices in Fuego and Scoop under different salt treatments. The enrichment index quantifies the relative accumulation or depletion of ions in the stomatal complex compared with the whole leaf. Salt treatments were coded as C (control), T1 (50 mM NaCl), T2 (100 mM NaCl), T3 (50 mM Na₂SO₄) and T4 (50 mM CaCl₂). Vectors indicate the contribution and direction of individual variables to the principal components. Points represent genotype positions in PCA space, and 95% confidence ellipses were constructed around genotype mean positions based on a multivariate t-distribution.

The PCA separated Fuego and Scoop mainly along PC1, reflecting distinct multivariate trait associations. Scoop was positioned predominantly on the positive side of PC1 and was associated with larger chloroplast surface area and volume, whereas Fuego was located toward the negative side of PC1 and showed stronger associations with Na depletion (more negative Na enrichment index) and higher SA:V ratio. The Cl enrichment index contributed positively to both PC1 and PC2, whereas photosynthetic rate (A) and chloroplast sphericity contributed negatively along PC2.

Treatment distribution further revealed genotype-specific responses under salinity. The Na₂SO₄ treatment produced the strongest separation between genotypes. Under this treatment, Fuego was characterized by lower SA:V ratio and stronger Na depletion, coinciding with pronounced chloroplast expansion and reduced photosynthetic performance observed in the corresponding analyses, whereas Scoop maintained a higher SA:V ratio with comparatively smaller chloroplasts. Control and NaCl treatments showed partial overlap between genotypes, while CaCl₂ and Na₂SO₄ resulted in greater divergence within the multivariate trait space.

The 95 % confidence ellipses confirmed clear separation between Fuego and Scoop, demonstrating distinct integrated responses involving chloroplast morphology, ion partitioning and photosynthetic performance under salt stress.

### 3.4 Genotype-specific relationships between ion enrichment, chloroplast morphology and photosynthetic performance

To determine the integrated relationships among photosynthetic performance, guard cell ion enrichment and chloroplast morphology, Pearson correlation matrices were constructed separately for Fuego and Scoop (Fig. 3). The two genotypes exhibited distinct correlation networks, indicating genotype-dependent associations between ion status, chloroplast remodeling and photosynthetic function.

**Figure 3.**
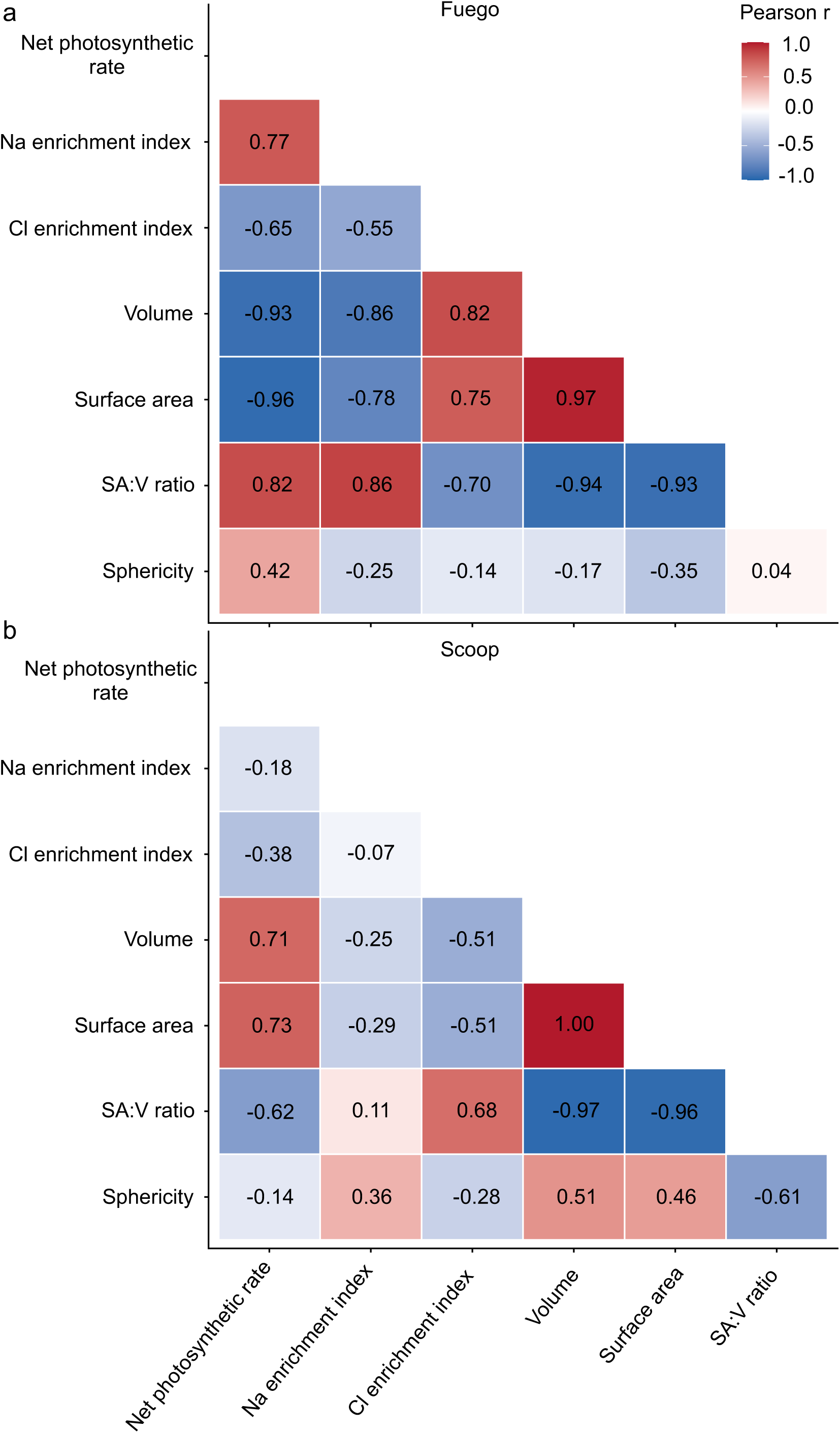
Correlation matrices of physiological and chloroplast morphological traits in faba bean genotypes. a,b, Pearson correlation matrices showing relationships among net photosynthetic rate, Na and Cl enrichment indices, and chloroplast morphological traits, including volume, surface area, surface area-to-volume ratio (SA:V) and sphericity, in Fuego (a) and Scoop (b). The enrichment index quantifies the relative accumulation or depletion of ions in the stomatal complex compared with the whole leaf. Color intensity indicates the strength and direction of correlations, with red representing positive and blue representing negative associations.

In Fuego (Fig. 3a), net photosynthetic rate (A) showed positive correlations with the Na enrichment index (r = 0.77) and the surface area-to-volume (SA:V) ratio (r = 0.82), but strong negative correlations with chloroplast volume (r = −0.93) and surface area (r = −0.96). Consistently, the Na enrichment index was positively associated with chloroplast volume (r = 0.82) and surface area (r = 0.75), whereas the Cl enrichment index showed negative relationships with chloroplast volume (r = −0.86) and surface area (r = −0.78). These associations indicate that chloroplast enlargement in Fuego was linked with reduced photosynthetic performance under salt stress.

In contrast, Scoop displayed an opposite relationship between chloroplast size and photosynthesis (Fig. 3b). A was positively correlated with chloroplast volume (r = 0.71) and surface area (r = 0.73), suggesting that larger chloroplast structures were associated with enhanced photosynthetic performance in this genotype. The Na enrichment index showed weak associations with A (r = −0.18) and chloroplast volume (r = −0.25), while the Cl enrichment index showed moderate negative correlations with A (r = −0.38), chloroplast volume (r = −0.51) and surface area (r = −0.51), together with a positive correlation with the SA:V ratio (r = 0.68).

Despite these genotype-specific responses, both Fuego and Scoop shared conserved chloroplast geometric relationships. Chloroplast volume and surface area were tightly correlated in both genotypes (r = 0.97 and 1.00, respectively), and both traits showed strong negative associations with the SA:V ratio (volume versus SA:V, r = −0.94 in Fuego and −0.97 in Scoop; surface area versus SA:V, r = −0.93 in Fuego and −0.96 in Scoop). Thus, although chloroplast geometry followed similar scaling relationships across genotypes, its association with photosynthetic performance differed markedly between Fuego and Scoop, highlighting genotype-specific functional consequences of chloroplast structural plasticity.

## 4 Discussion

Salt stress induces extensive changes in cellular structure and ion homeostasis (Haq et al., 2025; Zhao et al., 2020), but how these processes are integrated within guard cells to regulate physiological performance remains poorly understood. Chloroplasts are highly dynamic organelles that undergo structural and functional remodeling in response to environmental stress, contributing to adjustments in photosynthesis, metabolism and stress acclimation (Shu et al., 2013; Wang et al., 2024). Here, we demonstrate that guard cell chloroplast remodeling represents a genotype-dependent response to salinity that is closely associated with local ion partitioning and photosynthetic regulation. Rather than representing a passive consequence of stress injury, chloroplast structural plasticity appears to contribute to distinct adaptive strategies, revealing guard cell chloroplast architecture as an important functional component of salinity adaptation.

### 4.1 Genotype-specific guard cell chloroplast remodeling determines contrasting responses to salinity

Salt stress induced substantial but genotype-dependent remodeling of guard cell chloroplast architecture in faba bean (Fig. 1). Fuego showed pronounced chloroplast enlargement under stronger salinity, particularly under high NaCl and Na₂SO₄, accompanied by a lower surface area-to-volume (SA:V) ratio. In contrast, Scoop exhibited a more treatment-dependent structural response, characterized by smaller chloroplasts under high NaCl and CaCl₂ and higher SA:V ratios under Na₂SO₄ and CaCl₂. These contrasting patterns indicate that the two genotypes differ not only in the extent of chloroplast remodeling but also in how chloroplast architecture responds to the ionic environment under salinity.

Stress-induced chloroplast enlargement has been reported in a range of plant species and is commonly associated with ultrastructural reorganization. Electron microscopy studies have shown that salinity promotes starch granule accumulation, thylakoid swelling, grana distortion and changes in stromal volume, collectively contributing to chloroplast enlargement (Gao et al., 2015; Rahman et al., 2000; Shen et al., 2019). These changes are generally considered adaptive during the early stages of stress but may progress to chloroplast dysfunction under prolonged or severe salinity. Salt stress also frequently increases the number and size of starch granules within chloroplasts, further contributing to chloroplast expansion (Liang et al., 2024; Rahman et al., 2000; Shen et al., 2019; Wang et al., 2013). This response has been attributed to an imbalance between carbon assimilation and carbohydrate utilization, resulting in transient starch deposition during stress acclimation, although excessive starch accumulation may ultimately disrupt chloroplast ultrastructure and photosynthetic function (Rahman et al., 2000; Shen et al., 2019). Moreover, the co-localization of salt ions and starch granules within chloroplasts, revealed by advanced imaging approaches, suggests a close interaction between salt accumulation and carbon storage during salinity adaptation (Kanai et al., 2007; Noda et al., 2023). Therefore, the chloroplast enlargement observed in this study likely represents a coordinated response integrating salt adaptation with metabolic adjustment.

Importantly, the functional consequences of chloroplast remodeling differed between genotypes. In Fuego, increased chloroplast surface area and volume were strongly associated with reduced photosynthetic performance, suggesting that excessive structural expansion reflects stress-induced impairment rather than effective acclimation (Fig. 3a). In contrast, Scoop exhibited positive relationships between chloroplast size and photosynthetic rate, indicating that chloroplast enlargement may help sustain photosynthetic function under specific salt conditions (Fig. 3b). Thus, chloroplast size alone does not determine salt tolerance; rather, its physiological significance depends on the coordination between structural plasticity, elemental homeostasis and photosynthetic activity. These findings extend current understanding of stress-induced organelle plasticity by demonstrating that guard cell chloroplast architecture can follow distinct functional trajectories among genotypes under salinity.

### 4.2 Guard cell ion partitioning links chloroplast remodeling with photosynthetic regulation under salt stress

Beyond structural plasticity, our results indicate that guard cell chloroplast remodeling is closely associated with genotype-specific elemental partitioning under salinity. Fuego exhibited significantly stronger Na depletion from the stomatal complex than Scoop under NaCl stress, and this pattern coincided with contrasting chloroplast–photosynthesis relationships between the two genotypes (Table 1). In Fuego, greater Na enrichment was associated with chloroplast expansion and reduced photosynthetic performance, whereas Scoop showed weaker associations between Na enrichment, chloroplast morphology and photosynthetic activity (Fig. 3).

Element compartmentation is a central mechanism of plant salt tolerance because the distribution of Na and Cl among cellular compartments determines their effects on metabolism and photosynthetic function (Flowers et al., 2015; James et al., 2006; Mansour, 2023). Salt-tolerant plants minimize elemental toxicity by preferentially sequestering Na and Cl into vacuoles or specific cell types while maintaining low concentrations in metabolically active compartments, thereby preserving cellular homeostasis (Duan et al., 2023; Flowers et al., 2015; Teakle and Tyerman, 2010). Within the epidermis, elemental compartmentation across guard cells and surrounding epidermal cells is essential for maintaining osmotic balance and stomatal function. Stomatal movements depend on the coordinated regulation of K, Cl and other inorganic elements that generate guard-cell turgor, whereas excessive Na accumulation disrupts this balance, impairs stomatal behaviour and ultimately limits CO₂ assimilation (Jezek and Blatt, 2017; Lawson and Matthews, 2020; Zörb et al., 2022). Recent studies further demonstrate that salt-tolerant species maintain higher K/Na ratios and restrict excessive Na and Cl accumulation within guard cells, thereby preserving stomatal function under salinity (Franzisky et al., 2021; Karimi et al., 2021). Our findings extend these concepts by showing that elemental partitioning across the stomatal complex is closely linked to chloroplast remodeling and photosynthetic performance, suggesting that local elemental distribution determines whether chloroplast structural changes represent adaptive acclimation or stress-associated impairment under salinity.

Collectively, our results reveal contrasting genotype-specific associations among stomatal complex ion partitioning, chloroplast remodeling and photosynthetic performance under salinity (Fig. 4). Fuego showed stronger Na depletion, pronounced chloroplast expansion and reduced SA:V ratios, with larger chloroplasts associated with lower photosynthetic performance. In contrast, Scoop exhibited weaker Na depletion and more treatment-dependent chloroplast remodeling, with chloroplast size positively associated with photosynthetic performance. These contrasting patterns suggest that guard cell chloroplast architecture is closely linked to local elemental partitioning and may represent an important component of genotype-specific responses to salinity. More broadly, integrating subcellular ion profiling with 3D organelle phenotyping provides a spatially resolved framework for connecting elemental homeostasis with structural and physiological responses to salt stress.

**Figure 4.**
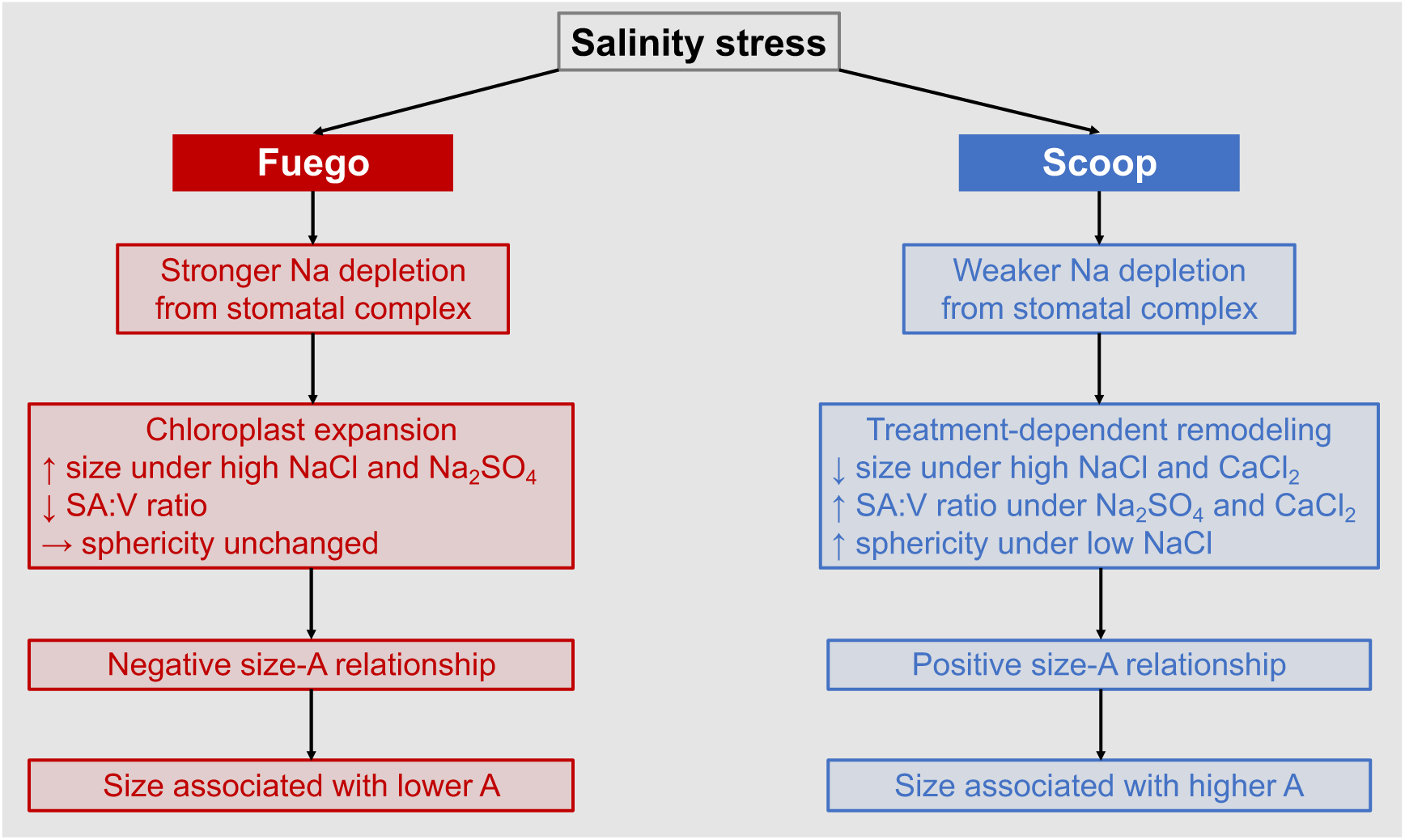
Conceptual model of genotype-specific guard cell responses to salinity. The model summarizes genotype-specific associations among Na partitioning, chloroplast remodeling, and photosynthetic performance in Fuego and Scoop under salt stress. A, net photosynthetic rate; size, chloroplast surface area and volume; SA:V ratio, surface area-to-volume ratio.

## 5 Conclusion

Our study reveals that guard cell chloroplast remodeling represents a dynamic and genotype-dependent component of salinity adaptation in faba bean. Salt stress altered chloroplast architecture in distinct ways between genotypes, and the functional consequences of these changes depended on their association with local ion partitioning and photosynthetic performance. By integrating chloroplast morphology, guard cell ion enrichment and photosynthetic responses, we demonstrate that chloroplast structural plasticity cannot be interpreted independently from the ionic environment of the stomatal complex. Instead, the coordination between ion homeostasis and chloroplast remodeling is associated with whether structural changes contribute to stress acclimation or physiological impairment. These findings identify guard cell chloroplast architecture as an additional functional trait linking subcellular organization with salt tolerance and provide new insight into the mechanisms underlying genotype-specific adaptation to salinity.

## Acknowledgement

The work was financially supported by the Deutsche Forschungsgemeinschaft (DFG, German Research Foundation, ZO 118/15–1), project number 491678431. Parts of the equipment used was supported by the EFRE EU fund (grant no. 2172959). We thank Dr. Claus J. Burkhardt and Dr. Birgit Schröppel at NMI Natural and Medical Sciences Institute, University of Tübingen for technical support with cryo-SEM-EDX system and data analysis. We also thank Ms. Christiane Beierle and Ms. Dagmar Repper for cryo-specimen preparation.

## Author Contributions

X.D.Z. led the conceptualization of the study, developed the methodology, performed the investigation, curated and analyzed the data, developed software and visualizations, administered the project, and wrote the original draft of the manuscript as well as the revised versions. G.H.W. contributed to the investigation by microscope-based experimental work. M.W. contributed to deep learning-based 3D modeling. C.Z. contributed to the conceptualization of the study, secured funding, administered the project, and reviewed and edited the manuscript. All authors read and approved the final manuscript and agree to be accountable for all aspects of the work, ensuring that questions related to the accuracy or integrity of any part of the research are appropriately addressed.

## AI use

During the preparation of this manuscript, AI-assisted technology was used solely to improve language clarity and readability. After using this tool/service, the authors reviewed and edited the content as needed and take full responsibility for the content of the published article.

## References

Collado-Arenal, A.M., Exposito-Rodriguez, M., Mullineaux, P.M., Olmedilla, A., Romero-Puertas, M.C., Sandalio, L.M., 2024. Cadmium exposure induced light/dark-and time-dependent redox changes at subcellular level in Arabidopsis plants. Journal of Hazardous Materials 477, 135164.

da Silva, W.A., Ferreira-Silva, M., Araújo, W.L., Nunes-Nesi, A., 2025. Guard cells and mesophyll: a delicate metabolic relationship. Trends in Plant Science 30(2), 125–127.

Dang, T., Piro, L., Pasini, C., Santelia, D., 2024. Starch metabolism in guard cells: at the intersection of environmental stimuli and stomatal movement. Plant Physiology 196(3), 1758–1777.

Duan, Y., Lei, T., Li, W., Jiang, M., Zhao, Z.a., Yu, X., Li, Y., Yang, L., Li, J., Gao, S., 2023. Enhanced Na^+^ and Cl^−^ sequestration and secretion selectivity contribute to high salt tolerance in the tetraploid recretohalophyte Plumbago auriculata Lam. Planta 257(3), 52.

Flowers, T.J., Munns, R., Colmer, T.D., 2015. Sodium chloride toxicity and the cellular basis of salt tolerance in halophytes. Annals of Botany 115(3), 419–431.

Franzisky, B.L., Geilfus, C.M., Romo-Pérez, M.L., Fehrle, I., Erban, A., Kopka, J., Zörb, C., 2021. Acclimatisation of guard cell metabolism to long-term salinity. Plant, Cell & Environment 44(3), 870–884.

Gao, H.-J., Yang, H.-Y., Bai, J.-P., Liang, X.-Y., Lou, Y., Zhang, J.-L., Wang, D., Zhang, J.-L., Niu, S.-Q., Chen, Y.-L., 2015. Ultrastructural and physiological responses of potato (Solanum tuberosum L.) plantlets to gradient saline stress. Frontiers in Plant Science 5, 787.

Granot, D., Kelly, G., 2019. Evolution of guard-cell theories: the story of sugars. Trends in Plant Science 24(6), 507–518.

Haq, S.I.U., Tariq, F., Sama, N.U., Jamal, H., Mohamed, H.I., 2025. Role of autophagy in plant growth and adaptation to salt stress. Planta 261(3), 49.

Hedrich, R., Shabala, S., 2018. Stomata in a saline world. Current Opinion in Plant Biology 46, 87–95.

James, R.A., Munns, R., Von Caemmerer, S., Trejo, C., Miller, C., Condon, T., 2006. Photosynthetic capacity is related to the cellular and subcellular partitioning of Na^+^, K^+^ and Cl^−^ in salt-affected barley and durum wheat. Plant, Cell & Environment 29(12), 2185–2197.

Jarvis, P., López-Juez, E., 2013. Biogenesis and homeostasis of chloroplasts and other plastids. Nature Reviews Molecular Cell Biology 14(12), 787–802.

Jezek, M., Blatt, M.R., 2017. The membrane transport system of the guard cell and its integration for stomatal dynamics. Plant Physiology 174(2), 487–519.

Kanai, M., Higuchi, K., Hagihara, T., Konishi, T., Ishii, T., Fujita, N., Nakamura, Y., Maeda, Y., Yoshiba, M., Tadano, T., 2007. Common reed produces starch granules at the shoot base in response to salt stress. New Phytologist 176(3), 572–580.

Karimi, S.M., Freund, M., Wager, B.M., Knoblauch, M., Fromm, J., M Mueller, H., Ache, P., Krischke, M., Mueller, M.J., Müller, T., 2021. Under salt stress guard cells rewire ion transport and abscisic acid signaling. New Phytologist 231(3), 1040–1055.

Kirchhoff, H., 2019. Chloroplast ultrastructure in plants. New Phytologist 223(2), 565–574.

Lambert, D., Peterson, B., Terpenning, I., 1991. Nondetects, detection limits, and the probability of detection. Journal of the American Statistical Association 86(414), 266–277.

Lawson, T., 2009. Guard cell photosynthesis and stomatal function. New Phytologist 181(1), 13–34.

Lawson, T., Matthews, J., 2020. Guard cell metabolism and stomatal function. Annual Review of Plant Biology 71(1), 273–302.

Liang, H., Shi, Q., Li, X., Gao, P., Feng, D., Zhang, X., Lu, Y., Yan, J., Shen, S., Zhao, J., 2024. Synergistic effects of carbon cycle metabolism and photosynthesis in Chinese cabbage under salt stress. Horticultural Plant Journal 10(2), 461–472.

Lim, S.L., Flütsch, S., Liu, J., Distefano, L., Santelia, D., Lim, B.L., 2022. Arabidopsis guard cell chloroplasts import cytosolic ATP for starch turnover and stomatal opening. Nature Communications 13(1), 652.

Mansour, M.M.F., 2023. Role of vacuolar membrane transport systems in plant salinity tolerance. Journal of Plant Growth Regulation 42(3), 1364–1401.

Martín-Fernández, J.A., Barceló-Vidal, C., Pawlowsky-Glahn, V., 2003. Dealing with zeros and missing values in compositional data sets using nonparametric imputation. Mathematical Geology 35(3), 253–278.

Munns, R., Tester, M., 2008. Mechanisms of salinity tolerance. Annual Review of Plant Biology 59, 651–681.

Naeem, M.S., Warusawitharana, H., Liu, H., Liu, D., Ahmad, R., Waraich, E.A., Xu, L., Zhou, W., 2012. 5-Aminolevulinic acid alleviates the salinity-induced changes in Brassica napus as revealed by the ultrastructural study of chloroplast. Plant Physiology and Biochemistry 57, 84–92.

Noda, Y., Hirose, A., Wakazaki, M., Sato, M., Toyooka, K., Kawachi, N., Furukawa, J., Tanoi, K., Naito, K., 2023. Starch-dependent sodium accumulation in the leaves of Vigna riukiuensis. Journal of Plant Research 136(5), 705–714.

Oi, T., Enomoto, S., Nakao, T., Arai, S., Yamane, K., Taniguchi, M., 2020. Three-dimensional ultrastructural change of chloroplasts in rice mesophyll cells responding to salt stress. Annals of Botany 125(5), 833–840.

Rahman, S., Matsumuro, T., Miyake, H., Takeoka, Y., 2000. Salinity-induced ultrastructural alterations in leaf cells of rice (Oryza sativa L.). Plant Production Science 3(4), 422–429.

Santelia, D., Lunn, J.E., 2017. Transitory starch metabolism in guard cells: unique features for a unique function. Plant Physiology 174(2), 539–549.

Schwenkert, S., Fernie, A.R., Geigenberger, P., Leister, D., Möhlmann, T., Naranjo, B., Neuhaus, H.E., 2022. Chloroplasts are key players to cope with light and temperature stress. Trends in Plant Science 27(6), 577–587.

Shen, J.-l., Wang, Y., Shu, S., Jahan, M.S., Zhong, M., Wu, J.-q., Sun, J., Guo, S.-r., 2019. Exogenous putrescine regulates leaf starch overaccumulation in cucumber under salt stress. Scientia Horticulturae 253, 99–110.

Shi, X., Hannon, N.M., Bloom, A.J., 2024. Metals and other ligands balance carbon fixation and photorespiration in chloroplasts. Physiologia Plantarum 176(4), e14463.

Shu, S., Yuan, L.-Y., Guo, S.-R., Sun, J., Yuan, Y.-H., 2013. Effects of exogenous spermine on chlorophyll fluorescence, antioxidant system and ultrastructure of chloroplasts in Cucumis sativus L. under salt stress. Plant Physiology and Biochemistry 63, 209–216.

Teakle, N.L., Tyerman, S.D., 2010. Mechanisms of Cl-transport contributing to salt tolerance. Plant, Cell & Environment 33(4), 566–589.

Thalmann, M., Santelia, D., 2017. Starch as a determinant of plant fitness under abiotic stress. New Phytologist 214(3), 943–951.

Wang, X., Chang, L., Wang, B., Wang, D., Li, P., Wang, L., Yi, X., Huang, Q., Peng, M., Guo, A., 2013. Comparative proteomics of Thellungiella halophila leaves from plants subjected to salinity reveals the importance of chloroplastic starch and soluble sugars in halophyte salt tolerance. Molecular & Cellular Proteomics 12(8), 2174–2195.

Wang, X., Chen, Z., Sui, N., 2024. Sensitivity and responses of chloroplasts to salt stress in plants. Frontiers in Plant Science 15, 1374086.

Yamane, K., Oi, T., Enomoto, S., Nakao, T., Arai, S., Miyake, H., Taniguchi, M., 2018. Three-dimensional ultrastructure of chloroplast pockets formed under salinity stress. Plant, Cell & Environment 41(3), 563–575.

Zeiger, E., Talbott, L.D., Frechilla, S., Srivastava, A., Zhu, J., 2002. The guard cell chloroplast: a perspective for the twenty-first century. New Phytologist 153(3), 415–424.

Zhao, C., Zhang, H., Song, C., Zhu, J., Shabala, S., 2020. Mechanisms of plant responses and adaptation to soil salinity. The innovation 1(1).

Zörb, C., Franzisky, B.L., Lehr, P.P., Kosch, R., Altenbuchinger, M., Geilfus, C.-M., 2022. Impact of nutritional imbalance on guard cell metabolism and stomata regulation under saline hyperosmotic conditions. Advances in Botanical Research 103, 123–138.

Zörb, C., Geilfus, C.M., Dietz, K.J., 2019. Salinity and crop yield. Plant Biology 21, 31–38.

