## supplementary files for "Guard Cell Chloroplast Remodeling Under Salt Stress Correlates with Ion Enrichment in Faba Bean"

**Supplementary Methods S1. Leaf ionomic analysis by ICP-OES and ion chromatography**

For bulk leaf ion analysis, approximately 50 mg of freeze-dried fifth leaf material was subjected to microwave-assisted pressure digestion with concentrated nitric acid (65% $HNO_3$) in quartz vessels using an UltraClave V microwave system (MLS, Leutkirch, Germany) according to established protocols (Wollmann et al., 2018). The digested samples were analysed by inductively coupled plasma–optical emission spectrometry (ICP-OES; Agilent 5110, Agilent Technologies, Santa Clara, CA, USA) for Na, K, Ca, Mg, P and S. Chloride (Cl) concentrations were determined separately by ion chromatography with suppressed conductivity detection.

Element concentrations were quantified using external calibration with certified multi-element and single-element standards. Samples with Na concentrations below the detection limit (<400 ppm) were assigned a value corresponding to half of the detection limit (200 ppm) for statistical analysis and visualization.

**Supplementary Methods S2. Cryo-SEM-EDX analysis of stomatal complex surface elemental composition**

Cryogenic scanning electron microscopy coupled with energy-dispersive X-ray spectroscopy (cryo-SEM-EDX) was used to characterize the relative surface elemental composition of stomatal complexes in faba bean leaves. This approach was applied to preserve the native distribution of mobile elements, particularly Na, K and Cl, while avoiding potential artefacts associated with chemical fixation, dehydration and embedding procedures that may introduce external ions or redistribute soluble elemental pools. Epidermal tissues were plunge-frozen in liquid nitrogen to preserve native elemental distributions (Franzisky et al., 2025).

Epidermal strips from the fifth fully expanded leaves used for gas exchange measurements were mounted on copper grids and transferred into a cryo-preparation chamber (Quorum Technologies) connected to a Zeiss Crossbeam 550 SEM. No freeze fracture or sublimation treatment was applied to minimize redistribution or loss of mobile ions. Samples were sputter-coated with platinum (5 mA, 45 s; approximately 35 nm) to reduce charging effects and improve X-ray signal stability.

SEM imaging and EDX measurements were performed at an accelerating voltage of 8 kV, beam current of 1 nA, magnification of 2126× and working distance of 5.5 mm. The sample stage was tilted by 45° to improve X-ray collection efficiency. Because stomatal complexes exhibit complex three-dimensional surface structures, cells with suitable orientation relative to the detector were manually selected to minimize shadowing effects and improve measurement consistency.

Elemental spectra were collected using an Oxford Instruments X-MaxN 150 SDD detector controlled by AZtec software (version 5.1). Spectral fitting included biological elements (C, N, O, Na, Mg, P, S, Cl, K and Ca) and instrumental signals originating from the copper support grid, platinum coating and gold sample holder. Quantification was performed using the AZtec standardless routine, including background subtraction, peak deconvolution, absorption correction, fluorescence correction and pulse pile-up correction.

Hydrogen cannot be detected by EDX. Carbon and oxygen were retained during normalization because they represent dominant structural components of plant tissues and provide a stable matrix reference for relative comparisons. Nitrogen was included during spectral fitting to improve peak separation but excluded from quantitative interpretation because the low-energy N Kα signal is strongly influenced by absorption, background interference and coating effects.

At the applied accelerating voltage of 8 kV, the electron interaction volume extended to approximately 1.5 μm in depth. Therefore, EDX signals primarily represent the elemental composition of the outer surface region of epidermal cells rather than intracellular ion concentrations. Accordingly, atom% values obtained by cryo-SEM-EDX were interpreted as relative surface elemental composition and not as absolute cellular ion contents. These measurements provide complementary spatial information to bulk leaf ion concentrations obtained by ICP-OES.

Element selection for biological interpretation was based on both spectral reliability and biological relevance. Although Na, Mg, P, S, Cl, K and Ca were detectable under the applied conditions, Na, K and Cl were selected for comparative analysis because they showed clear characteristic peaks and consistent detection across genotype–treatment combinations. Representative EDX spectra from all genotype–treatment combinations are shown in Supplementary Figure S1.

Regions of interest corresponding to the entire stomatal complex were manually defined according to SEM morphology. The stomatal complex was analysed as a single functional unit without separating individual component cells. At least 10 stomatal complexes per treatment were analysed from epidermal strips collected from five independent leaf samples. Each spectrum was acquired for at least 5 min. Elemental abundances were expressed as atomic percentages (atom%) and normalized to the sum of reliably detected biological elements (C, O, Na, Mg, P, S, Cl, K and Ca) for semi-quantitative comparison.

$$Atom\%_{\,normalized} = \frac{Atom\%_{element}}{\sum(C + O + Na + Mg + P + S + Cl + K + Ca)} \times 100$$

When elemental signals, including low Na or Cl signals, were indistinguishable from background noise and lacked clear characteristic peaks, values were considered below the detection limit and assigned as zero for statistical analysis and visualization. This criterion enabled consistent comparison of relative elemental distribution patterns among genotypes and salt treatments.

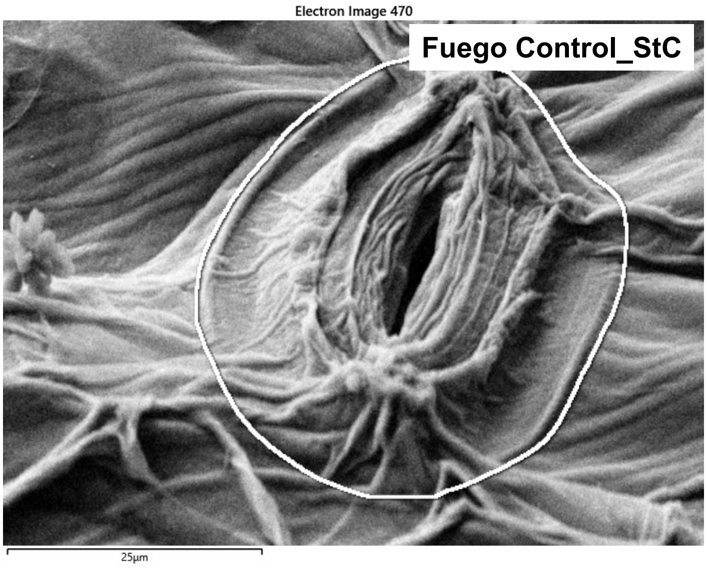
Electron Image 470
Fuego Control_StC
25µm

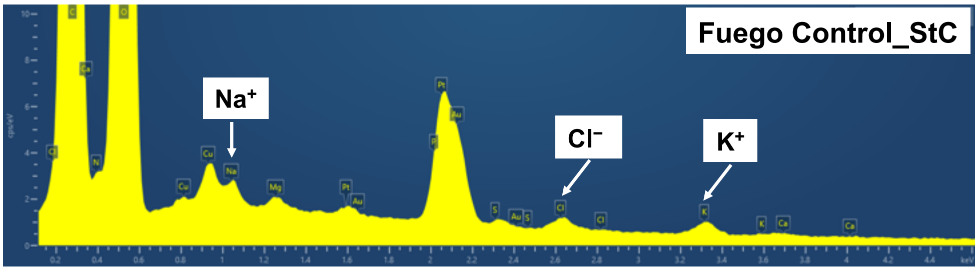
Fuego Control_StC
Na+
Cl−
K+

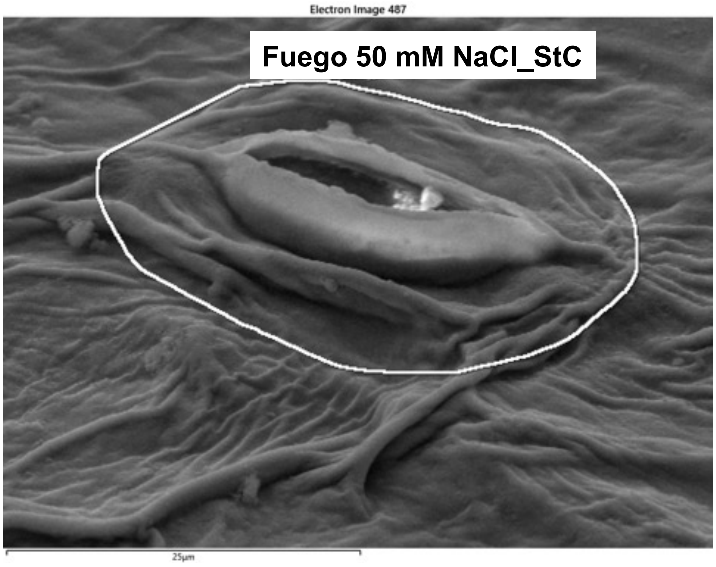
Electron Image 487
Fuego 50 mM NaCl_StC
25µm

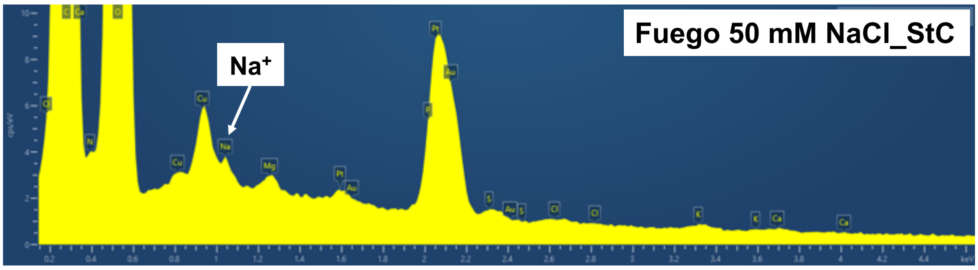
Fuego 50 mM NaCl_StC
Na+

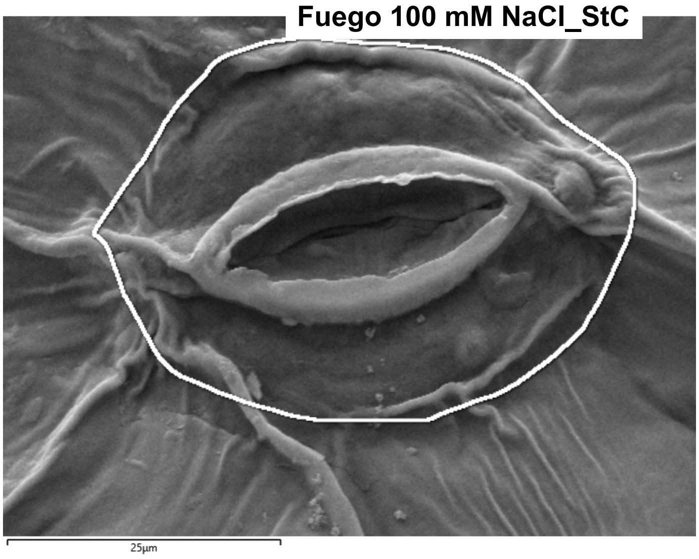
Fuego 100 mM NaCl_StC
25µm

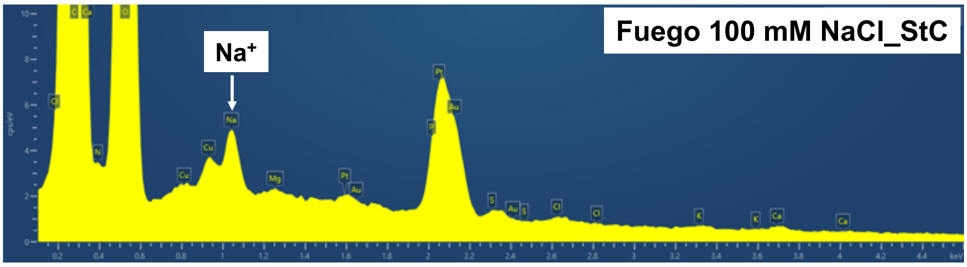
Fuego 100 mM NaCl_StC
Na+

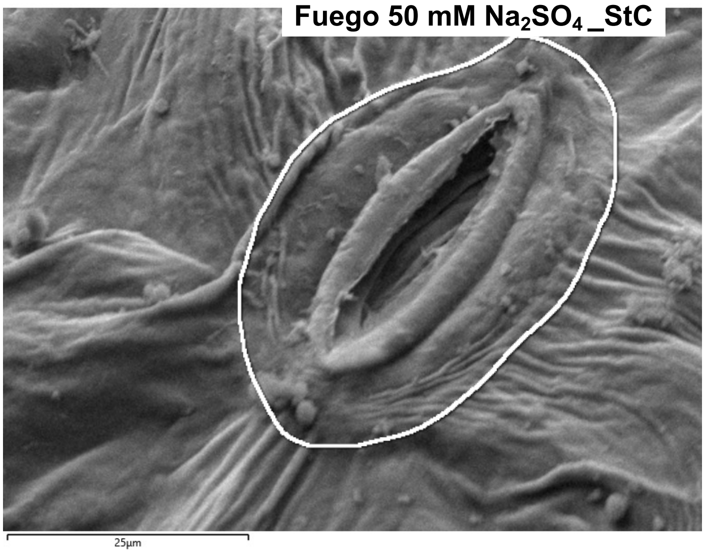
Fuego 50 mM Na2SO4_StC
25µm

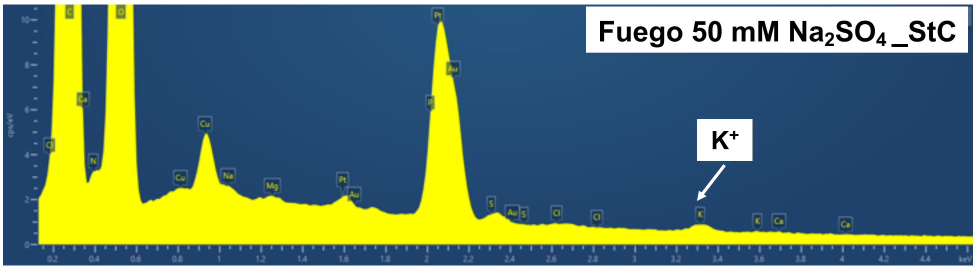
Fuego 50 mM Na2SO4_StC
K+

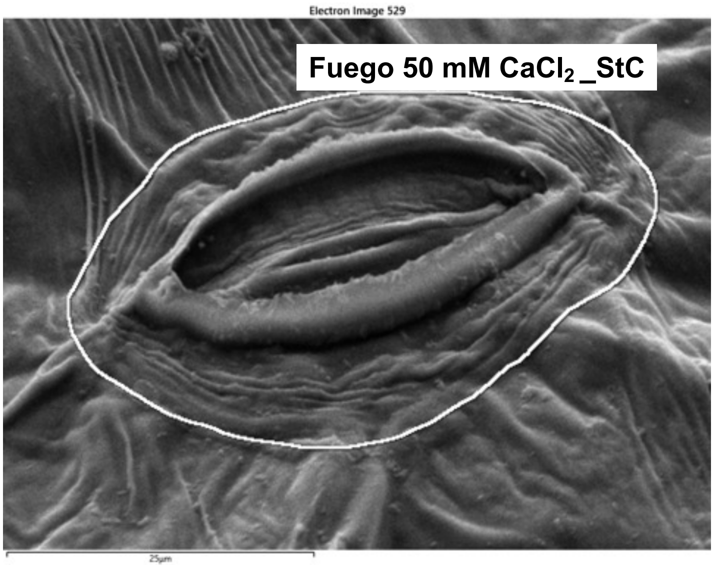
Electron Image 529
Fuego 50 mM $CaCl_2$_StC
25µm

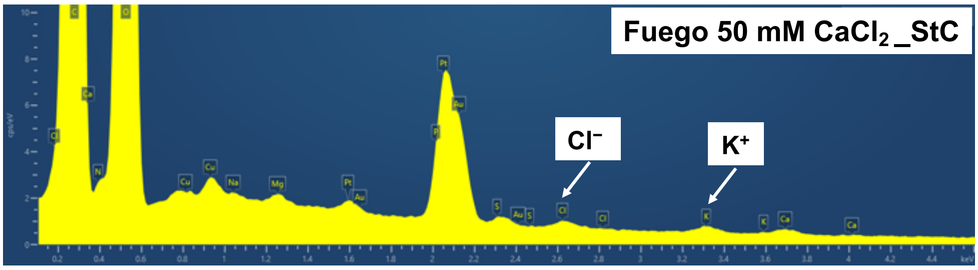
Fuego 50 mM $CaCl_2$_StC
$Cl^-$
$K^+$

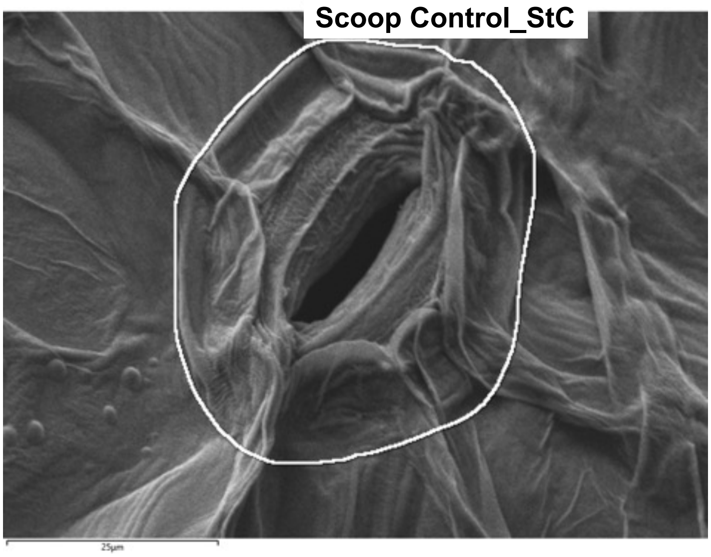
Scoop Control_StC
25µm

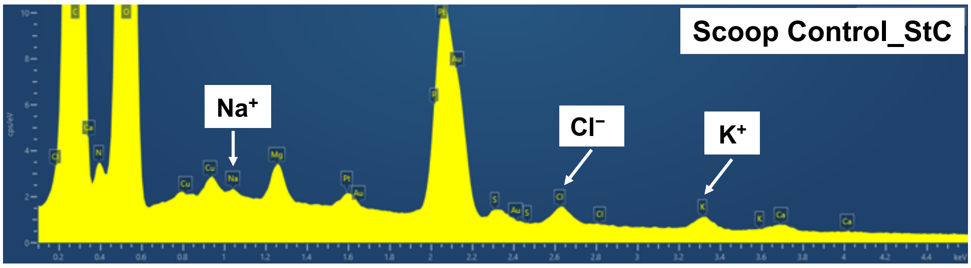
Scoop Control_StC
$Na^+$
$Cl^-$
$K^+$

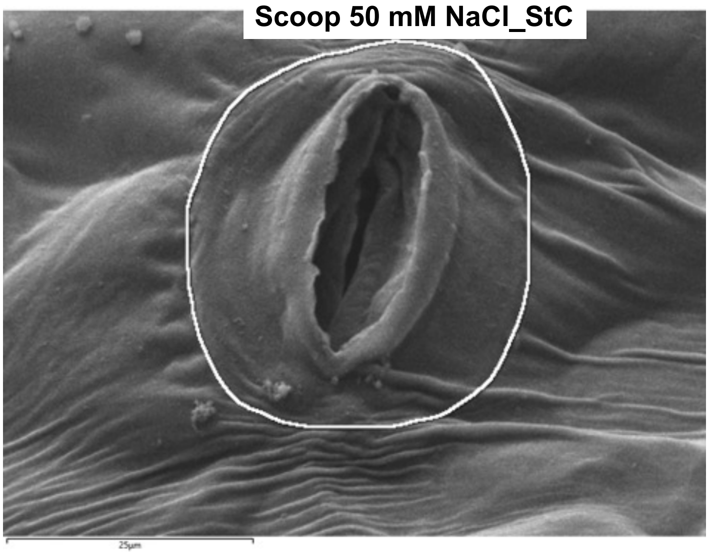
Scoop 50 mM NaCl_StC
25μm

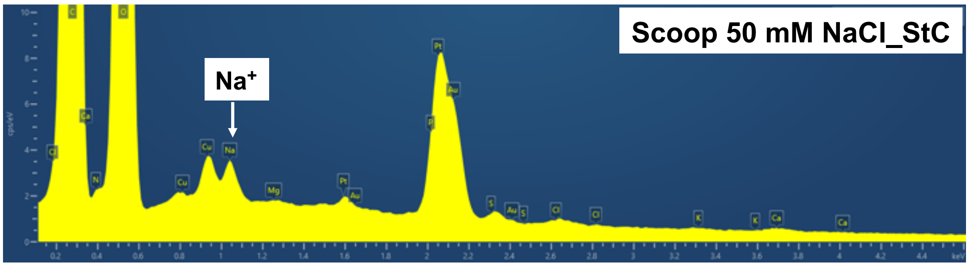
Scoop 50 mM NaCl_StC
$Na^+$

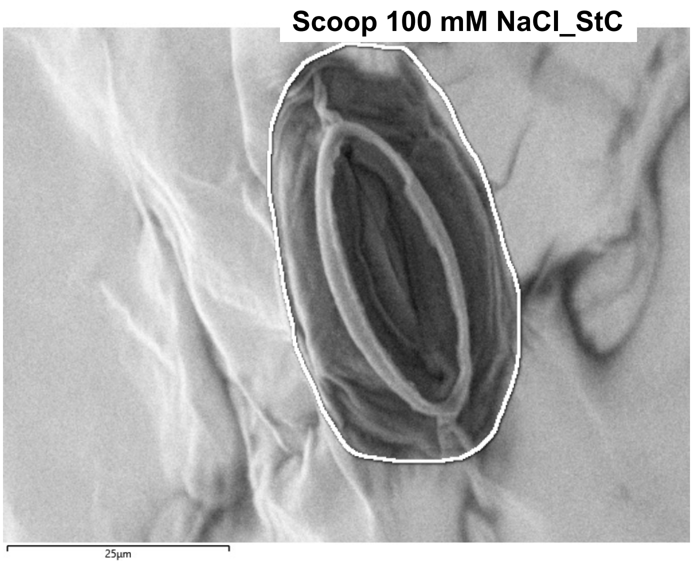
Scoop 100 mM NaCl_StC
25μm

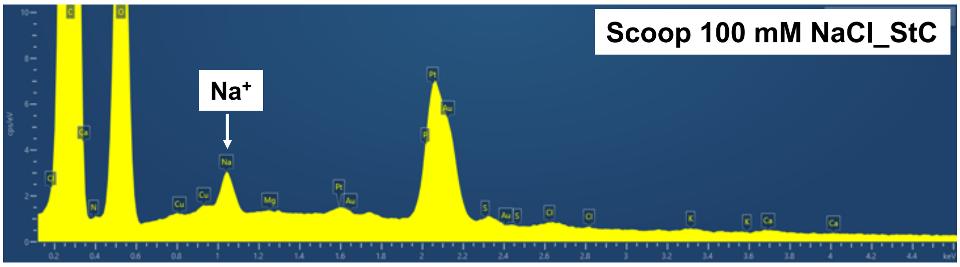
Scoop 100 mM NaCl_StC
$Na^+$

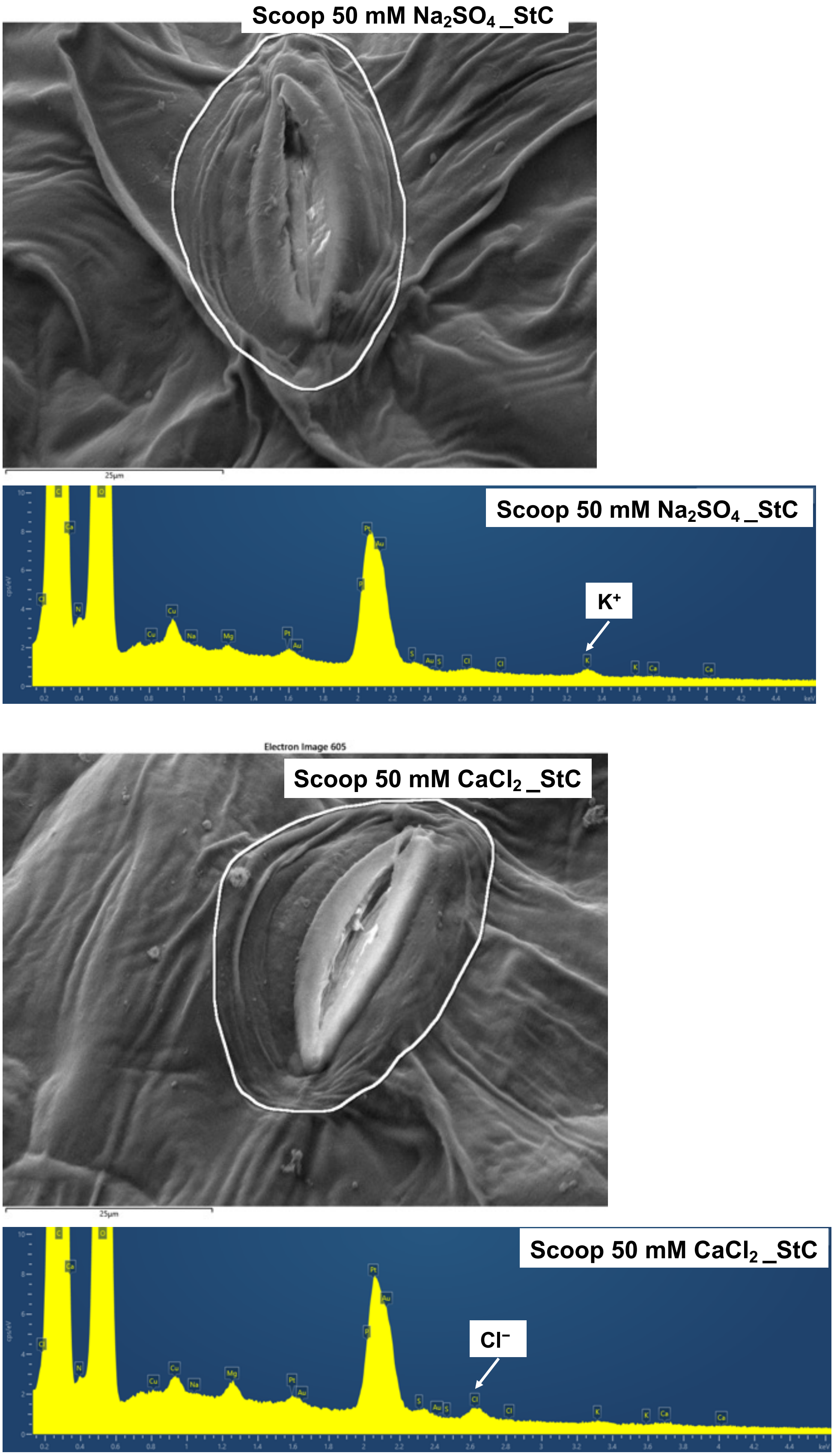


**Supplementary Figure S1. Representative cryo-SEM-EDX spectra of stomatal complex surface regions across faba bean genotypes and salt treatments**

Representative energy-dispersive X-ray spectroscopy (EDX) spectra obtained from stomatal complex surface regions of Fuego and Scoop under control and salt treatments. Spectra from all genotype–treatment combinations are shown to demonstrate elemental detection reliability across experimental conditions. Characteristic peaks corresponding to Na, K and Cl are specifically labelled because these elements were selected for comparative analysis of salt-induced elemental distribution patterns. Other detected elements, including Mg and Ca, were retained during spectral fitting and normalization and are shown only through automatic peak annotation in the exported spectra. Signals originating from the copper support grid (Cu), platinum coating (Pt) and gold sample holder (Au) are also automatically annotated. EDX signals represent relative elemental composition of the near-surface region, with a detection depth of approximately 1.5 μm, rather than intracellular ion concentrations.

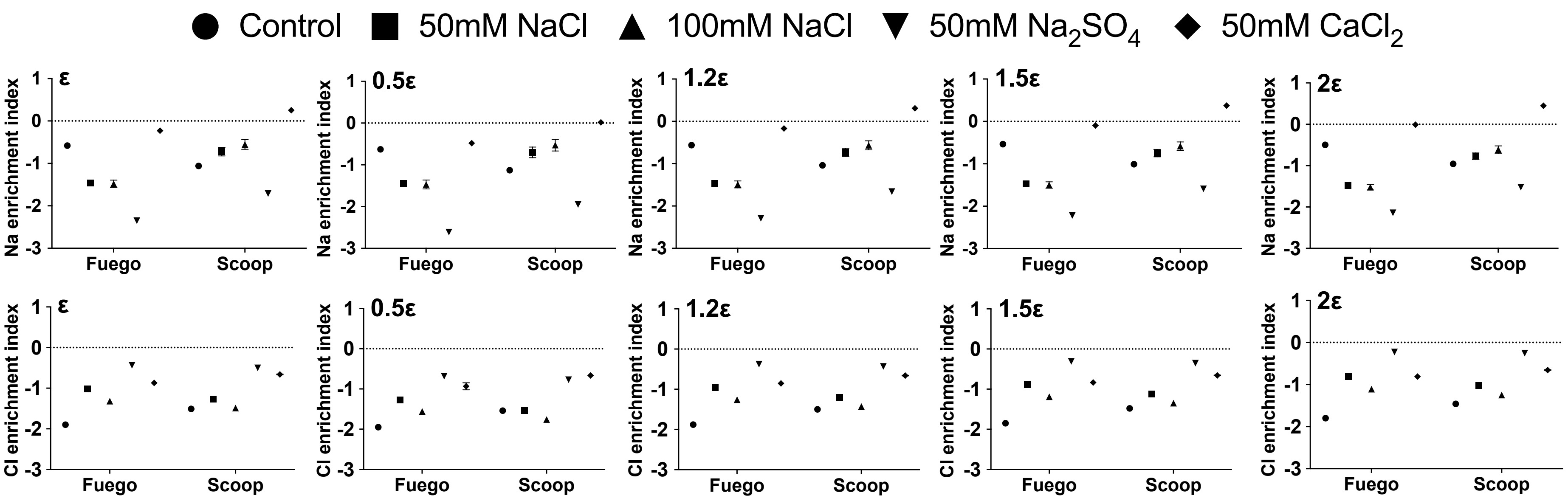


**Supplementary Figure S2. Sensitivity analysis of pseudocount selection for Na and Cl enrichment indices in two faba bean genotypes**

Sensitivity analysis showing the effect of different pseudocount values (ε) on Na and Cl enrichment indices calculated from stomatal complex surface elemental signals relative to corresponding whole-leaf ion concentrations in two faba bean genotypes, Fuego and Scoop. The reference ε value was defined as half of the minimum non-zero value across all samples, and alternative values corresponding to 0.5×, 1×, 1.2×, 1.5× and 2× of this value were evaluated. Similar trends in treatment- and genotype-associated enrichment patterns were observed across the tested ε values, supporting the selection of the reference pseudocount for enrichment index calculation. The enrichment index was calculated as the $\log_{10}$ ratio between normalized stomatal complex surface elemental signals and corresponding normalized whole-leaf ion concentrations.

**Supplementary Table S1. Evaluation of chloroplast volume thresholds for quality control of segmented chloroplast objects**

| Threshold ($\mu m^3$) | Removed objects | Removed (%) | Interpretation |
|---|---|---|---|
| 0.01 | 8 | 0.71 | Minimal filtering |
| 0.05 | 51 | 4.53 | Limited artifact removal |
| 0.1 | 65 | 5.78 | Moderate filtering |
| 0.2 | 80 | 7.11 | Selected QC threshold |
| 0.5 | 92 | 8.18 | Increasing risk of excluding biological objects |
| 1 | 116 | 10.31 | Potential over-filtering |

The raw dataset contained 1,125 segmented chloroplast objects obtained from three-dimensional confocal reconstructions. Different minimum volume thresholds were evaluated to identify an appropriate quality-control cutoff for removal of potential segmentation artifacts. A threshold of 0.2 $\mu m^3$ was selected because it excluded 80 small chloroplast objects (7.11% of detected objects) while retaining 1,045 objects (>90% of the original dataset), thereby minimizing potential loss of biologically relevant chloroplast structures. All downstream analyses were performed using chloroplast objects with volumes $\geq 0.2$ $\mu m^3$.

**References**

Franzisky, B.L., Zhang, X., Burkhardt, C.J., Majorovits, E., Hummel, E., Schertel, A., Geilfus, C.-M., Zörb, C., 2025. Application of cryo-FIB-SEM for investigating ultrastructure in guard cells of higher plants. Plant Physiology and Biochemistry 220, 109546.

Wollmann, I., Müller, T., Breuer, J., Möller, K., Augustenberg, L.T., 2018. Standardisiertes Verfahren zur Bewertung des Phosphordüngewerts von Recyclingdüngemitteln. Ministerium für Umwelt, Klima und Energiewirtschaft, Referat 25.
